# Optogenetic Kindling Model Reveals Functional Synaptopathy as a Driver of Temporal Lobe Epilepsy and Associated Comorbidities

**DOI:** 10.64898/2026.11.22.768446

**Authors:** Hyunyoung Kim, Na-Young Seo, Woojae Butterwort, Jae-Won Jang, Ji-young Kim, Sohee Kim, Kea Joo Lee, Hyungju Park

## Abstract

Temporal lobe epilepsy (TLE) is one of the most prevalent and pharmaco-resistant forms of focal epilepsy in adults. The precise cellular and network mechanisms underlying TLE progression remain elusive. Animal models are essential for studying molecular and cellular mechanisms of TLE; however, widely used models largely rely on acute seizure induction and fail to recapitulate the gradual, progressive course of human TLE. Here, we present an optogenetic kindling mouse model of TLE, in which repeated stimulation induces a progressive increase in seizure susceptibility. This mouse model exhibits prominent TLE phenotypes, including robust hippocampal epileptiform activity, behavioral seizures, emotional changes, reactive astrogliosis, and impaired hippocampal learning and memory. Moreover, hippocampal synaptic analyses of this model revealed altered synaptic transmission and plasticity, without structural changes in excitatory or inhibitory synapses or overt cell death. These findings demonstrate that progressive epileptogenesis, synaptic dysfunction and TLE-associated phenotypes can develop independently of structural remodeling or overt cell death. This model provides a platform for investigating the synaptic mechanisms underlying TLE-associated synaptopathy.

## Introduction

Epilepsy is a neurological disorder affecting approximately 50 million individuals worldwide (Feigin et al., 2025). Spontaneous, recurrent seizures accompanied by a broad spectrum of neuropsychiatric and cognitive comorbidities, including anxiety, depression, and memory impairment, are well established clinical phenotypes of epilepsy. Particularly, TLE represents one of the most prevalent and pharmacoresistant forms of epilepsy in adults (Téllez-Zenteno and Hernández-Ronquillo, 2012). While anti-seizure medications provide symptomatic relief for about one-third of patients, most patients remain refractory to current treatments (Chen et al., 2018). Therefore, defining the pathophysiological mechanisms underlying chronic epileptogenesis is urgently needed.

Despite decades of research, the cellular and circuit mechanisms underlying TLE progression remain unclear. This gap stems largely from limitations of existing animal models. For example, chemical convulsant models, typically induced by systemic or intra-hippocampal administrations of pilocarpine or kainic acid, have been the mainstay of TLE research (Lévesque et al., 2016). While successful for evoking immediate, excessive, and uncontrolled seizure activity (Lévesque et al., 2016), these models are inadequate for studying features of TLE that are important for understanding its etiology, such as gradual and chronic disease progression. Moreover, widespread necrotic brain damage or systemic drug toxicity complicates the isolation of primary pathophysiological mechanisms of TLE.

Epileptogenesis, the progressive transition from a normal state to one of heightened susceptibility to recurrent seizures, has long been associated with structural and functional reorganization of hippocampal circuits (Buckmaster and Dudek, 1997; Curia et al., 2008; Goldberg and Coulter, 2013). Classical models, such as chemoconvulsant-induced TLE, have revealed prominent synaptic remodeling including mossy fiber sprouting, granule cell dispersion, and loss of inhibitory interneurons (Buckmaster and Dudek, 1997; Curia et al., 2008; Lévesque and Avoli, 2013; Mello et al., 1993; Wuarin and Dudek, 1996). However, these changes are typically accompanied by extensive neuronal death and neuroinflammatory responses such as reactive gliosis, which arise from the severe, acute status epilepticus used to trigger epileptogenesis (Lévesque and Avoli, 2013; Mello et al., 1993; Patel et al., 2019; Thom, 2009; Vezzani et al., 2022). Importantly, this pattern of severe neuronal loss does not reflect all human TLE: a substantial fraction of patients with TLE show no hippocampal sclerosis on MRI (Vaughan et al., 2016; Wang et al., 2013), and histopathological classification of resected hippocampi includes a distinct category characterized by gliosis without the characteristic pattern of neuronal loss (Blümcke et al., 2013). Thus, in existing models, the cellular alterations and inflammatory responses confound the interpretation of whether circuit remodeling is a primary driver of epileptogenesis or a secondary consequence of neurodegeneration and inflammation.

To address this, we established an optogenetic kindling model by repeatedly stimulating excitatory neurons in the ventral CA1 (vCA1) of the hippocampus (Tescarollo et al., 2023), a region implicated in the initiation of spontaneous seizures in TLE (Buckmaster et al., 2022).

Our optogenetic TLE model recapitulated characteristics of TLE including epileptiform activity and clinically relevant behavioral phenotypes. Notably, these features co-occurred with synaptic dysfunction that manifested as altered spontaneous and evoked synaptic transmission and impaired long-term synaptic plasticity, in the absence of structural synapse remodeling or cell death. Furthermore, this functional impairment was accompanied by the upregulation of transmembrane protein 230 (TMEM230), an excitatory synapse-specific vesicular protein overexpressed in human TLE patients (Zhang et al., 2024), validating the clinical relevance of our model. Altogether, these findings demonstrate that TLE pathogenesis can be driven primarily by functional synaptopathy and establish an experimental platform for investigating TLE etiology.

## Materials and Method

All mouse experiments were performed in accordance with all ethical regulations and institutional guidelines approved by the Institutional Animal Care and Use Committee (IACUC) of the KBRI (IACUC-24-00020-M1).

### Mice

The following mouse lines were used: CamKIIα-CreERT2 mice (The Jackson Laboratory) and ExN-ChR2 (CamKIIα-CreERT2:Ai32) mice. All the mouse lines were maintained on a C57BL/6 genetic background. All mice were housed in a specific pathogen-free room at the Laboratory Animal Center in the KBRI with feed (ENVIGO) and water. The room was kept at 20–24 °C and 40–60% humidity under a 12–12 h light–dark cycle with dark hours between 20:00 and 08:00. To induce ChR2 expression, tamoxifen (75 mg/kg) was intraperitoneally injected for 5 consecutive days. All mice were individually housed after implantation of optic fiber. The mice for behavioral, electrophysiological, and histological analyses were 9-12 weeks-old male and the mouse used for LFP recordings was 10 weeks-old female.

### Implantation of optic fiber

The mice were anesthetized with isoflurane (Hana pharm) and implanted with an optic fiber cannula (0.39 NA, 200 mm diameter) for laser stimulations (Thorlabs). The tip of the optic fiber cannula was positioned at unilateral vCA1 (AP, −3.10 mm; ML, 3.70 mm; DV, −2.5 mm). After surgery, the animals were individually housed in a clean cage to recover.

### Optrode preparation

A custom-made optrode was constructed by attaching a neural electrode array to an optic fiber cannula (Supplementary Fig. 1). Neural electrode arrays were fabricated using polyimide as the flexible substrate and encapsulation layer, with Cr/Au (25/200 nm) serving as the conductive traces and recording sites. The device features a 10.5-mm-long and 450-μm-wide shank with 16 contacts, incorporating microelectrode sites with an opening area of 40 × 40 μm² at a pitch of 100 μm. Electrochemical impedance spectroscopy (EIS) showed an average impedance of 808.36 ± 482.38 kΩ at 1 kHz for the fabricated electrodes. The fabricated electrode array was connected to a printed circuit board (PCB) via anisotropic conductive film (ACF) bonding, and a 16-channel connector (Omnetics) was soldered onto the PCB. The PCB-electrode assembly and an optical fiber were integrated into a 3D-printed housing, where the electrode shank and the optical fiber were bonded together using polyethylene glycol (PEG).

### Local field potential recording

Light stimulation (LS) and local field potential (LFP) recordings were performed simultaneously using an optrode in which the optic fiber and the recording electrode were integrated into a single device. The optrode was implanted into the unilateral vCA1 at the same coordinates used for optic fiber implantation, and a contralateral S1 (AP, +1.50 mm, ML, -3.00 mm) bone screw anchored in the skull served as the reference and ground. LFPs were acquired with an RHS stimulation/recording system (Intan Technologies) at a sampling rate of 30 kHz and analyzed using MATLAB (MathWorks). For each LS session, LFPs were recorded from 5 min before to 5 min after LS, and a 2-min window consisting of 1 min before and 1 min after LS onset was used for analysis.

To visualize the temporal changes in oscillatory activity before, during, and after LS, spectrograms were computed with the continuous wavelet transform using an analytic Morlet wavelet, and wavelet power was expressed in dB from 1 to 150 Hz. To quantify these changes in each frequency band, PSD was then compared between the pre-stimulus baseline and stimulation epochs. The 60-s segment immediately before the first light pulse of each LS unit was used as the baseline, and the 3-s segments corresponding to each of the five light bursts were used as stimulation epochs. Mean PSD was calculated in the delta (0.5-4 Hz), theta (4-8 Hz), alpha (8-12 Hz), beta (12-30 Hz), gamma (30-80 Hz), and high-gamma (80-150 Hz) bands, averaged across the five bursts, and expressed relative to baseline as 10·log10 (PLS/baseline) (dB).

For inter-spike interval (ISI) analysis, epileptiform spikes were detected as positive deflections exceeding five times the standard deviation (SD) of the baseline signal. For each threshold crossing, the time of the local peak was taken as the spike time, and peaks separated by less than 50 ms were treated as a single spike. ISIs were calculated as the time differences between consecutive spikes, and their mean and SD were obtained for each LS session.

### Behavioral experiments

Behavioral testing started the day after the last day of opogenetic kindling. Prior to the test, mice were habituated to the testing room for 1 hour.

#### Optogenetic kindling

The protocol consisted of three phases: initial kindling, the first re-kindling, and the second re-kindling. One week after optic fiber cannula implantation, optogenetic kindling was initiated with daily laser stimulation over five consecutive days. Each daily session consisted of six stimulation units delivered at 30-minute intervals. A single stimulation unit comprised five light bursts delivered every 3 seconds (total duration: ∼30 seconds per unit). For each light pulse, blue light (470 nm, Thorlabs) was delivered at 50 Hz with a 25% duty cycle and an irradiance of 2 mW at the fiber tip. Two subsequent re-kindling sessions were conducted at two-week intervals following the initial kindling. Each re-kindling was conducted on a single day (days 28 and 42; Fig. 1b), using the same stimulation parameters as the initial kindling (six stimulation units at 30-minute intervals; Fig. 1c). The mice were video-recorded for 5 minutes starting from the onset of the first light pulse in each stimulation unit. Seizure severity was scored from recorded video using a modified Racine scale (RS) adapted previous study(Tescarollo et al., 2023). RS was categorized as follows: RS0, normal behaviors; RS1, arrest or sudden motion; RS2, chewing and/or head nodding; RS3, forelimb clonus; RS4, fore- and hindlimb clonus with rearing; RS5, loss of posture and/or falling; RS6, brief seizure behavior; and RS7, severe seizure behavior with jumping or squeaking. A termination criterion was applied across all kindling and re-kindling sessions: if an animal displayed seizure behaviors rated RS5 or higher for three consecutive stimulation units, the session was immediately discontinued. For initial kindling, any remaining daily sessions were omitted once this criterion was met. Similarly, re-kindling sessions were terminated upon reaching the same behavioral threshold.

**Figure 1.**
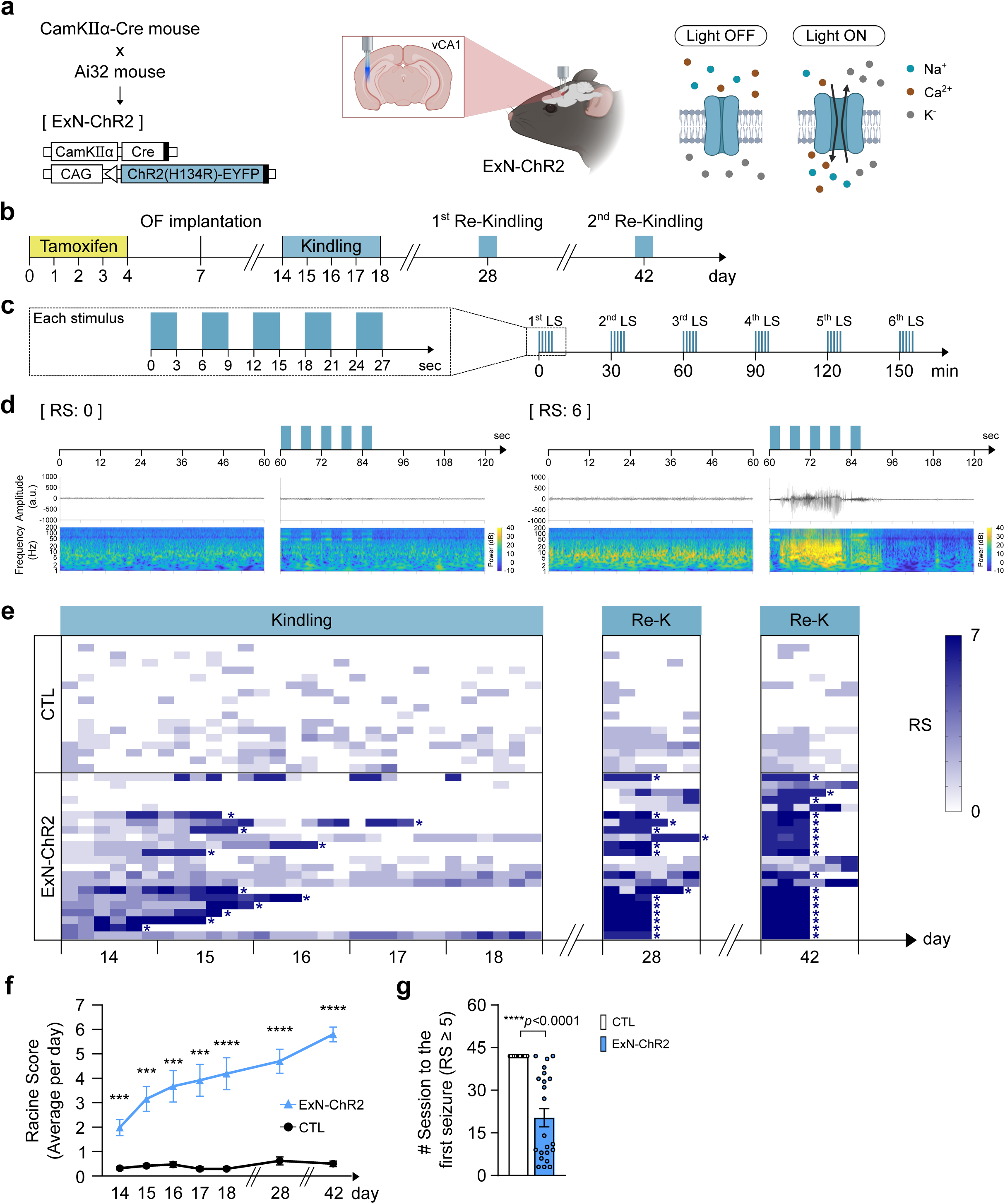
Establishment of an optogenetic mouse model of temporal lobe epilepsy. a. Schematic of the ExN-ChR2 line and optogenetic stimulation. This schematic illustration was made using Biorender. b. Experimental timeline. Tamoxifen injection (days 0–4), optic fiber (OF) implantation (day 7), initial optogenetic kindling (days 14–18), the first re-kindling (day 28), and the second re-kindling (day 42). c. Illustration showing kindling protocol per day. d. Representative trace of vCA1 LFP recording from a mouse showing no seizure response (Racine score, RS 0; left) or a generalized seizure (RS 6; right). For each, the light train patterns (top), corresponding LFP traces (middle) and spectrogram (bottom) are shown. Blue bars, light stimulation (LS). e. Heat map summary of the RS of individual mice for every LS session. Each row is one mouse and each column one LS session. Color bar, RS 0–7. Asterisk indicates the termination of kindling due to kindling criterion protocol. CTL, n = 18 mice; ExN-ChR2, n = 22 mice. f. Summary of averaged RS per day. ***p = 0.00039 (day 14); ***p = 0.00017 (day 15); ***p = 0.00046 (day 16); ***p = 0.00011 (day 17); ****p = 0.000043 (day 18); ****p = 0.00000021 (day 28); ****p < 1 × 10^-13^ (day 42); two-way ANOVA with Sidak’s post hoc test. g. Number of LS sessions required to reach the first generalized seizure (RS ≥ 5). ****p = 0.0000004, unpaired two-tailed Student’s t-test. All data are presented as mean ± SEM.

#### Open field test

Open field test was used to determine general locomotion and anxiety levels. The assessment took place in a square arena within a specially designed sound attenuating chamber. The square arena was 30 cm × 30 cm × 25 cm (Length × Width × Height). The mouse was placed in the center of the arena and allowed to freely explore the arena for 5 min while being tracked by an automated tracking system of SMART3.0 (Panlab) software. Distance moved and time spent in predefined zones of the arena were measured. The arena was cleaned with distilled water and 70% ethanol at the end of the trial.

#### Forced swimming test

Forced swimming test was performed in a plastic cylinder (diameter: 11 cm, height: 26 cm) filled to a depth of 18.5 cm with water (22 ± 1). The mice were placed in the water with their heads above the surface while being recorded by a video camera, and their immobility time during the first 5 min was manually quantified.

#### Contextual fear conditioning test

Contextual fear conditioning tests were performed in sound-attenuating chambers equipped with a grid floor and a movement-detection sensor (Panlab). Freezing behavior was automatically measured and analyzed using Packwin software (Panlab). During the fear-conditioning phase, the mice were habituated to the shock chamber for 2 min and then subjected to five-foot shocks (0.3 mA, 0.5 seconds) with a 1-min-interval. The mice remained in the chamber for an additional 1 min after the shock and then returned to their home cages. Twenty-four hours later, contextual retrieval testing was performed for 3 min in the same chamber, during which freezing time was automatically recorded by the floor sensor.

### Electrophysiology

All chemicals for the standard artificial cerebral spinal fluid (ACSF) were purchased from Sigma unless indicated.

### Brain slice preparation and extracellular field recordings

For extracellular field excitatory postsynaptic potential (fEPSP) recordings, acute hippocampal slices were prepared in a modified sucrose-based slicing ACSF (in mM: 5 KCl, 1.25 NaH_2_PO_4_, 26 NaHCO_3_, 0.5 CaCl_2_, 10 MgCl_2_, 10 D-glucose, and 212.5 sucrose). Transverse hippocampal slices (400 μm thick) were prepared using a VF-200-OZ Compresstome (Precisionary) in ice-cold slicing ACSF and recovered at 27 °C for 50 min in standard ACSF (in mM: 124 NaCl, 5 KCl, 1.25 NaH_2_PO_4_, 26 NaHCO_3_, 10 D-glucose, 2.5 CaCl_2_, and 1.5 MgCl_2_) before recording. Slices were placed in a recording chamber and continuously perfused (2-3 ml/min) with oxygenated standard ACSF at 27 °C. fEPSPs were recorded through low-impedance (2.5-3.5 MΩ) glass micropipettes filled with standard ACSF placed in the CA1 SR. Test stimuli were applied at 0.05 Hz. Stimulus intensity was set by producing an I/O relationship of fEPSP slopes against stimulus intensities. For LTP recording, stimulus intensity was adjusted to 50% of the maximum fEPSP slope. After a stable 10-min baseline, LTP was induced by five trains of theta burst stimulation (TBS; each train consisting of four stimuli at 100 Hz, with 200-ms intervals between trains) delivered through a tungsten bipolar electrode placed in the SR. fEPSP slopes were analyzed using WinLTP software. TBS-induced short-term potentiation (STP), reflecting a post-tetanic potentiation-like component, was calculated by averaging fEPSP slopes over the first 5 min after LTP induction, expressed as a percentage of the baseline average. The maintenance of LTP was evaluated by averaging fEPSP slopes during the last 50-60 min after LTP induction.

### Whole-cell patch clamp

For whole-cell patch clamp recordings, acute brain slices were obtained by coronal sectioning. Mice were anesthetized with isoflurane and transcardially perfused with ∼20 ml of the slicing ACSF (in mM: 93 N-methyl-D-glucamine (NMDG)-Cl, 2.5 KCl, 1.2 NaH_2_PO_4_, 30 NaHCO_3_, 20 HEPES, 5 sodium ascorbate, 2 Thiourea, 3 sodium pyruvate, 12 N-acetyl-L-cysteine (NAc), 0.5 CaCl_2_, 10 MgCl_2_, and 25 glucose; pH 7.3; 300-310 mOsm) before the brain was dissected. The brain was coronally sectioned at 400-µm thick using a VF-200-OZ Compresstome (Precisionary) in the slicing ACSF. Brain slices were recovered at 32.5 °C in the recovery ACSF (in mM: 104 NaCl, 2.5 KCl, 1.2 NaH_2_PO_4_, 24 NaHCO_3_, 5 HEPES, 5 sodium ascorbate, 2 Thiourea, 3 sodium pyruvate, 12 NAc, 2 CaCl2, 2 MgCl_2_, and 13 glucose; pH 7.3; 300-310 mOsm) for 1 h before recording. Slices were then placed in continuously perfused (2–3 ml/min) with the oxygenated standard ACSF (in mM: 124 NaCl, 2.5 KCl, 1.2 NaH_2_PO_4_, 24 NaHCO_3_, 5 HEPES, 2 CaCl_2_, 2 MgCl_2_, and 13 glucose; pH 7.3; 300-310 mOsm). To evoke EPSCs and IPSCs, a bipolar electrode (CBBPE75; FHC) was placed in CA1 SR region (Supplementary Fig. 3).

Data were acquired through a Multiclamp 700B amplifier (Molecular Devices), filtered at 5 kHz, digitized at 10–50 kHz, stored on a computer, and analyzed offline using pCLAMP 10 software (Axon Instruments). Borosilicate glass patch electrodes with a resistance of 3–5 MΩ filled with pipette solution containing (in mM) 140 Cs-methanesulfonate (Cs-MeSO_4_), 7 NaCl, 2 MgCl_2_, 4 Mg-ATP, 0.3 Na_3_-GTP, 10 HEPES, and 1 mM QX-314 (pH 7.3, 290–300 mOsm). All whole-cell patch clamp recording data were analyzed using pClamp 11.4.1.

### Immunohistochemistry

The mice were anesthetized and transcardially perfused with phosphate-buffered saline (PBS), followed by 4% paraformaldehyde (PFA, Biosesang) following perfusion with. Brain tissues were post-fixed in 4% PFA overnight subsequently transferred to a 30% sucrose solution in PBS for 48 h prior to sectioning. Serial coronal sections of the brain (thickness, 40 µm) were collected using a cryostat (Leica, CM3050S) and washed with PBS three times.

Brain sections were blocked with a blocking solution containing 10% normal goat or donkey serum and 0.2% Triton X-100 in PBS for 1 hour at room temperature, followed by overnight incubation at 4°C with primary antibody solution. For TMEM230 immunostaining, brain sections were incubated in pre-heated citrate buffer (pH 6.0) at 95°C for 10 min with shaking for antigen retrieval. Then, sections were allowed to cool at room temperature for 20 min. After washing three times with PBS, blocking was subsequently performed. The primary antibodies used for immunostaining were a rabbit anti-NeuN (1:1000, Abcam), a guinea pig anti-Vglut1 (1:2000, Millipore Sigma), a rabbit anti-Homer1 (1:500, Synaptic Systems), a guinea pig anti-VGAT (1:500, Synaptic Systems), a rabbit anti-Gephyrin (1:500, Synaptic Systems), a chicken anti-GFAP (1:500, Aves Labs), a guinea piganti-S100β (1:500, Synaptic Systems), a rabbit anti-Iba1 (1:500, Fujifilm Wako), a rabbit anti-TMEM230 (1:100, Proteintech), and the fluorescent secondary antibodies were diluted with blocking solution at a concentration of 1:250 and incubated with the sections for 2 h at room temperature. The fluorescent secondary antibodies used were a goat anti-guinea pig IgG Alexa Fluor 568-conjugated antibody (Thermo Scientific), anti-rabbit IgG Alexa Fluor 555-conjugated antibody (Thermo Scientific), a goat anti-rabbit IgG Alexa Fluor 647-conjugated antibody (Thermo Scientific), and a donkey anti-chicken IgY Alexa Fluor 594-conjugated antibody (Thermo Scientific). Nuclei were fluorescently labeled with DAPI containing mounting medium (Vectashield).

#### Neuronal loss analysis

To measure the thickness of the CA1 pyramidal layer, confocal images of the stratum pyramidale (SP) were acquired using a Leica STELLARIS 8 confocal microscope equipped with a 40x water immersion objective lens and LAS X software. Four confocal images per mouse were analyzed, and their mean values were used for statistical comparison (n= 1 per mouse).

#### Synapse density

To quantify synapse density, confocal images of the hippocampal CA1 stratum radiatum (SR) were acquired under the same imaging condition described above. To assess excitatory and inhibitory synapses, puncta numbers of presynaptic markers (Vglut1 for excitatory, VGAT for inhibitory), postsynaptic markers (Homer1 for excitatory, Gephyrin for inhibitory), and their corresponding colocalized (double-positive) puncta were counted using the ImageJ (NIH, v1.54p) DIANA plugin. All puncta densities were normalized to an area of 100 μm^2^ or 10 μm^2^. Eight confocal images per mouse were analyzed, and their mean values were used for statistical comparison (n= 1 per mouse).

#### Sholl analysis

To evaluate the morphological complexity and gliosis of astrocyte and microglia, Sholl analyses were performed using maximum-intensity projection images generated from confocal z-stack images acquired using either a Leica STELLARIS 8 or a Andor Dragonfly 502w confocal microscope. Images were thresholded, converted to binary image, and skeletonized using ImageJ (NIH, v1.54p). The center of each cell soma was selected as the origin, from which concentric circles were generated at radial increments of 4 μm for GFAP and 2.5 μm for S100β and Iba1. The number of intersections was measured using the Sholl Analysis plugin and total length of process was measured using Analyze skeleton plugin in ImageJ. 6 to 10 cells per mouse were analyzed, and their mean values were used for statistical comparison (n= 1 per mouse).

#### TMEM230 expression

To examine TMEM230 expression level, the fluorescence intensity from eight confocal images per mouse was analyzed, and their mean value was used as a single biological replicate (n= 1 per mouse).

## Statistics

All the statistical analyses were performed via GraphPad Prism 11 with 95% confidence, and all the values are expressed as the mean ± standard error of the mean (SEM). Unpaired two-tailed Student’s t tests or Welch’s t tests were used to compare two groups. For multiple comparisons over time or distance, two-way ANOVA with Sidak’s post hoc tests was used. P-values less than 0.05 were considered statistically significant. All experiments were repeated at least 3 times independently.

## Results

### Generation of optogenetic TLE model

As human TLE frequently originates from focal hippocampal hyperactivity(Vossler et al., 2004), we designed an experimental strategy to mimic the origination of hippocampal hyperactivity by selectively stimulating the excitatory pyramidal neurons within the hippocampal CA1 region (Tescarollo et al., 2023). The most efficient and precise kindling of excitatory neurons is light-induced stimulation of channelrhodopsin-2 (ChR2)-expressing pyramidal neurons in the hippocampus. To achieve this, we cross-bred CamKIIα-CreERT2 mice with Ai32 reporter (*loxP-stop-loxP-ChR2-EYFP* knock-in line) mice to generate ExN-ChR2 mice (excitatory neurons expressing ChR2, Fig. 1a; see the Materials and Methods for detailed information). Tamoxifen-administered CamKIIα-CreERT2 mice lacking the Ai32 allele were used as control (CTL). An optical fiber was implanted into the vCA1 to target the principal seizure-onset zone characteristic of human TLE (Buckmaster et al., 2022).

We first evaluated whether focal hyperactivation of vCA1 excitatory neurons leads to epileptiform activity during optogenetic kindling sessions by monitoring changes in local field potentials (LFPs) in the hippocampal CA1 region. Our LFP recording results showed that repeated LS successfully induced a robust epileptiform activity throughout the kindling process (Fig. 1b-d; Supplementary Fig. 1, and 2a, e). Quantification of the power spectral density (PSD) further revealed a broadband increase in network oscillations during light-stimulation compared to baseline. During sessions 1-5 of the first re-kindling, spectral power increases were restricted to high-frequency bands (Supplementary Fig. 2d), consistent with the early emergence of high-frequency activity as a biomarker of epileptogenesis (Bragin et al., 2010). In contrast, during session 6 of the first re-kindling and throughout the second re-kindling, spectral power increased across all frequency bands (Supplementary Fig. 2d and 2h), which likely reflects the broadband contribution of frequent interictal spikes (Jabran et al., 2020).

**Figure 2.**
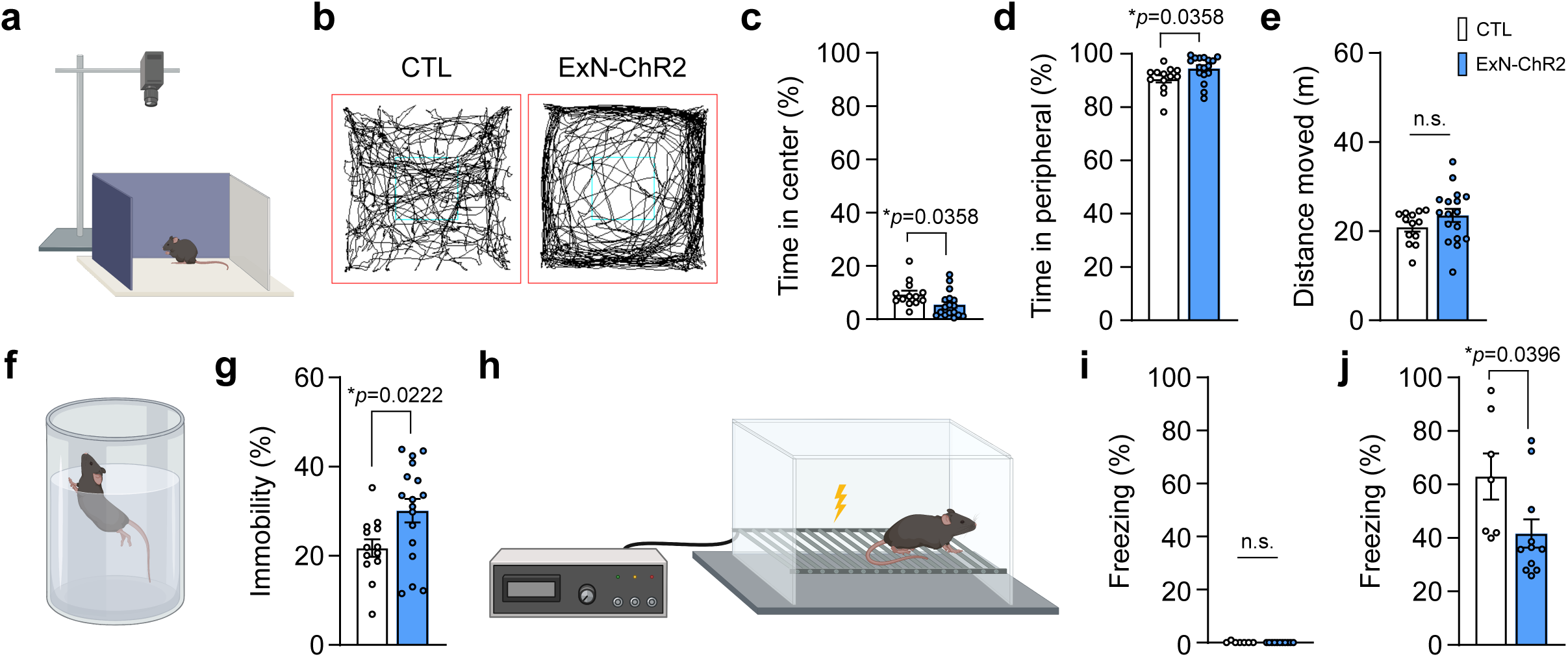
Behavioral comorbidities and cognitive deficits in optogenetic epilepsy mice. a-e. Open field test (OFT). CTL: n = 13, ExN-ChR2 n = 17. a. Schematic of the OFT. b. Representative traces of mice movement. Time spent in center (c; *p = 0.0358) and peripheral (d; *p = 0.0358). e. Total distance moved. P > 0.05. f-g. Forced swim test (FST). CTL: n = 13, ExN-ChR2 n = 17. f. Illustration of the FST. g. Time of immobility (%). *p = 0.0222. h-j. Contextual fear conditioning test (CFCT). CTL: n = 7, ExN-ChR2 n= 11. h. Schematic diagram of the CFCT. Time of freezing of baseline (i; p > 0.05) and retrieval (j, *p = 0.0396). All comparisons were tested using unpaired two-tailed Student’s t-test. All data are presented as mean ± SEM. All schematic illustrations in this figure were made using Biorender.

Next, we monitored changes in behavioral seizure phenotypes upon optogenetic kindling. Behavioral seizure scores were measured using the modified Racine scale (RS) by scoring visible behavioral outcomes during optogenetic stimulation relative to pre-stimulation baseline (see the Materials and Methods). Our analysis results showed that RS gradually transits from short arrest to generalized tonic seizures (Fig. 1e-g). During the initial kindling phase, 50% of mice were defined as fully kindled because they achieved the kindling criterion (RS ≥ 5 for three consecutive stimulation units). This success rate progressively increased to 64% and 77% during the first and second re-kindling sessions, respectively. Although a subset of mice did not meet the kindling criterion, all mice (100%) displayed at least one seizure event during the second re-kindling phase (Fig. 1e). The daily mean RS of ExN-ChR2 mice exhibited a steady upward linear trend over time and were significantly higher than those of CTL mice across all kindling phases (Fig. 1f), indicating an escalation of seizure severity. CTL mice lacking ChR2 showed no epileptiform LFP activity or behavioral seizures under identical light-stimulation (Fig. 1e-g), confirming that the observed seizures were induced by ChR2-mediated neuronal activation rather than LS per se.

### Behavioral deficits induced by optogenetic epileptogenesis

Patients with epilepsy, particularly those with TLE, are heavily burdened by comorbid psychiatric disorders and cognitive impairment (Fiest et al., 2013; Hermann et al., 2006; Keezer et al., 2016; Vinti et al., 2021). We next tested whether optogenetic kindling of ExN-ChR2 mice reproduces clinical comorbidities, such as anxiety, depression-like behaviors, and hippocampal cognitive deficits.

In the open field test (OFT; Fig. 2a, b), kindled ExN-ChR2 mice showed a significant reduction in the time spent in the center compared to CTL mice (Fig. 2c, d), without alteration in total distance moved (Fig. 2e), indicating heightened anxiety-like behavior. In the forced swim test (FST; Fig. 2f), kindled ExN-ChR2 mice exhibited a marked increase in immobility time, reflecting a depression-like state (Fig. 2g). Assessment of hippocampus-dependent associative memory using the contextual fear conditioning test (CFCT; Fig. 2h) showed that kindled ExN-ChR2 mice demonstrated significantly decreased contextual memory twenty-four hours post-conditioning (Fig. 2j), indicating attenuated hippocampal learning and memory.

Collectively, these findings indicate that chronic focal hyperactivity drives both seizure activity and the psychiatric and cognitive comorbidities of clinical TLE.

### Dysregulation of synaptic plasticity and transmission by optogenetic epileptogenesis

To elucidate the cellular substrates responsible for the observed network hyperexcitability and behavioral deficits, we performed *ex vivo* whole-cell patch-clamp and field potential recordings in acute hippocampal slices. First, we tested whether activity-dependent circuit plasticity was altered in kindled ExN-ChR2 mice by examining long-term potentiation (LTP) at the Schaffer-collateral-CA1 pathway. Hippocampal LTP was significantly reduced in slices from the kindled ExN-ChR2 mice (Fig. 3a-d), providing a cellular correlate for the attenuated hippocampal memory (Fig. 2j). Given the intact basal synaptic responses represented by the input/output (I/O) curve of field excitatory postsynaptic potentials (fEPSPs) across varying stimulus intensities (Fig. 3e-f), such LTP reduction in kindled ExN-ChR2 appears not to be caused by reduced synaptic strength.

**Figure 3.**
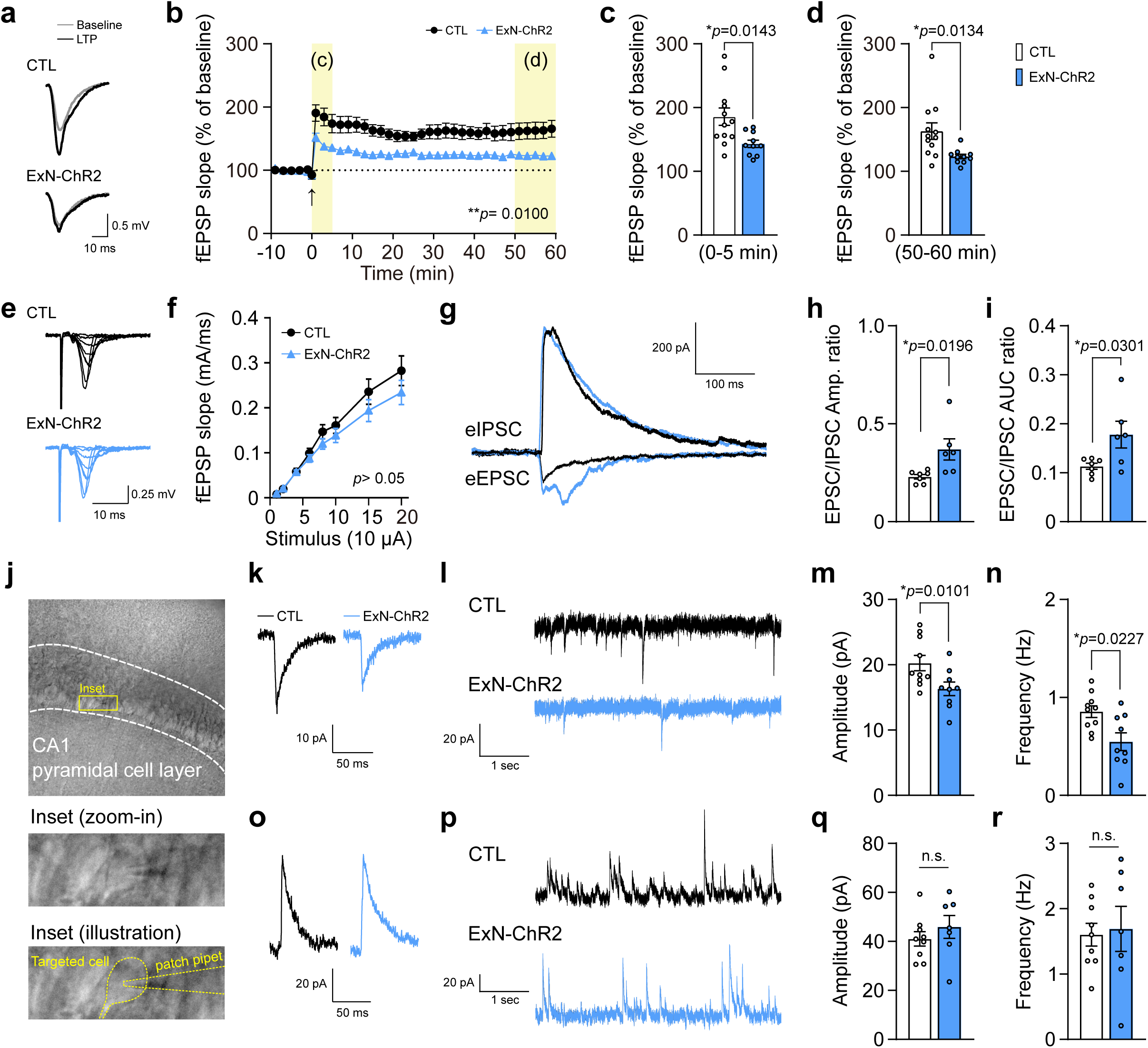
Reduced long-term potentiation and altered excitation/inhibition balance in CA1 pyramidal neurons of kindled ExN-ChR2 mice. a. Representative field EPSP (fEPSP) traces measured at the Schaffer-collateral pathway before (baseline, grey) and 50–60 min after LTP induction (black). b. Summary of average percentages of fEPSP slope normalized to baseline. Theta-burst stimulation was delivered at t = 0 min (arrow). CTL, n = 10 slices from 3 mice; ExN-ChR2, n = 12 slices from 3 mice. **p = 0.0100, two-way ANOVA. c-d. Average percentages of fEPSP slope during 0–5 min (c) and 50–60 min (d) after LTP induction. *p = 0.0143 (c), *p = 0.0134 (d), unpaired two-tailed Student’s t-test. e-f. fEPSPs evoked by increasing stimulus intensity. e. Representative traces of fEPSP in response to various ranges of stimulation intensities (10 ∼ 200 μA). f. Summary of input–output relationships curves for basal synaptic strength. p> 0.05, two-way ANOVA. g–i. Representative traces of evoked EPSCs (eEPSCs) and IPSCs (eIPSCs). CTL, n = 7 cells from 3 mice; ExN-ChR2, n = 6 cells from 5 mice. h. Average ratio of eEPSC/eIPSC peak amplitudes. *p = 0.0196, unpaired two-tailed Student’s t-test. i. Averaged ratio of eEPSC/eIPSC charge transfer estimated from area under curve (AUC) of eEPSCs and eIPSCs. *p = 0.0301, unpaired two-tailed Student’s t-test. j. Representative images showing whole-cell patch clamp recording of CA1 pyramidal neuron. The CA1 pyramidal cell layer was outlined (top, dashed lines). Yellow box, inset region (middle) showing recorded neuron. The same inset with the targeted cell and patch pipette outlined (bottom) was demonstrated. k. Representative averaged sEPSC responses. l. Representative sEPSC recording trace for 5 seconds. m-n. Average sEPSC amplitudes (m; *p = 0.0101) and frequency (n; *p = 0.0227). CTL, n = 10 cells from 7 mice, ExN-ChR2, n = 9 cells from 4 mice. Unpaired two-tailed Student’s t-test. o. Representative averaged sIPSC responses. p. Representative sIPSC recording trace for 5 seconds. q-r. Average sIPSC amplitudes (q; p > 0.05) and frequency (r; p > 0.05). CTL, n = 9 cells from 5 mice; ExN-ChR2, n = 7 cells from 5 mice. Unpaired two-tailed Student’s t-test. All data are presented as mean ± SEM.

Further analysis of changes in excitatory and inhibitory synaptic transmission revealed abnormal excitation-inhibition (E/I) balance (Supplementary Fig. 3). We found that both the ratios of peak amplitude and the Area Under the Curve (AUC) of evoked EPSC/IPSCs were significantly elevated in ExN-ChR2 mice (Fig. 3g-i), indicating a striking shift toward hyperexcitability of the hippocampal circuit. This imbalance of E/I ratio suggests that repetitive epileptic and synchronized circuit activity has evoked the mechanisms underlying hyperexcitable circuit activity and led to the increased anxiety and depression-like behaviors(Li et al., 2023; Tao et al., 2024).

Intriguingly, slices from ExN-ChR2 mice displayed a significant decrease in both the amplitude and frequency of spontaneous EPSCs (sEPSCs), whereas spontaneous IPSCs (sIPSCs) remained unchanged (Fig. 3k-r). Given no significant changes in membrane properties of CA1 neurons (Table 1), the reduced sEPSC amplitude and frequency are unlikely to be caused by altered neuronal properties. Together with the elevated evoked E/I ratio (Fig. 3g-i), these findings indicate a complex dysregulation of synaptic transmission, possibly involving a homeostatic downregulation of spontaneous excitatory transmission.

**Table 1.**
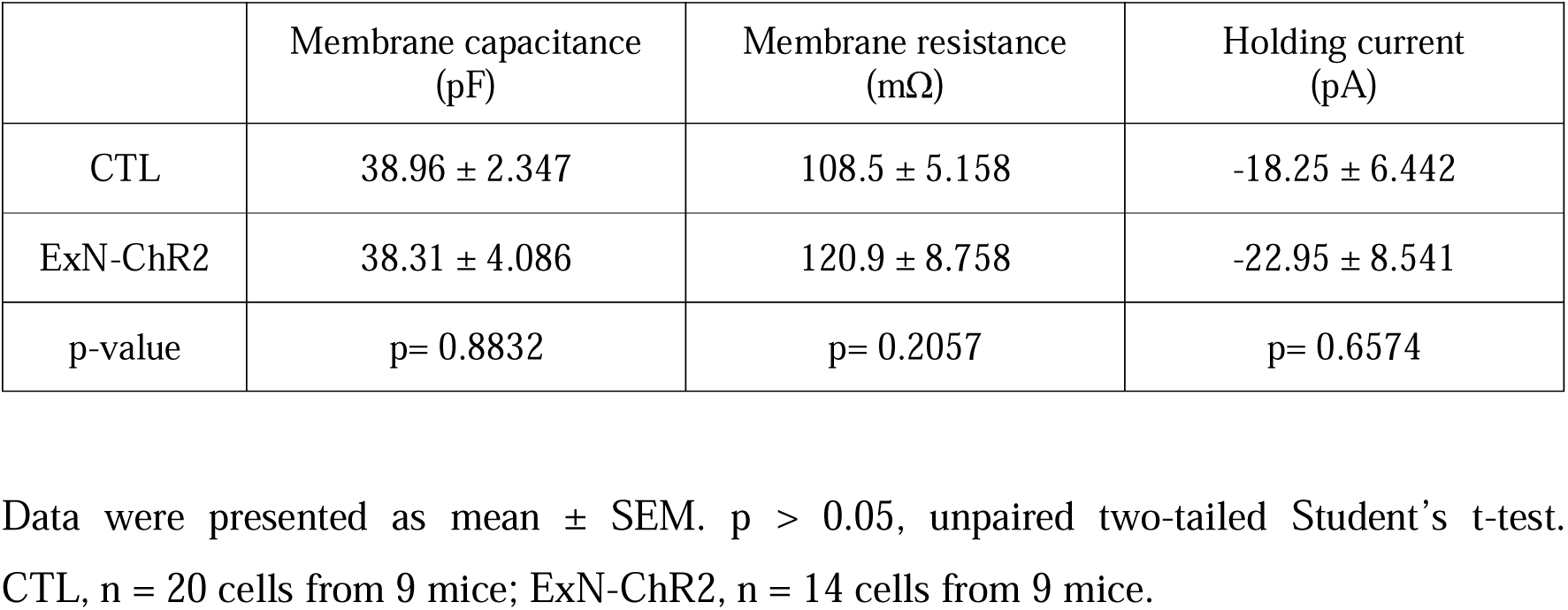
Electrophysiological properties of recorded CA1 pyramidal neurons.

|  | Membrane capacitance<br>(pF) | Membrane resistance<br>(mΩ) | Holding current<br>(pA) |
|---|---|---|---|
| CTL | 38.96 ± 2.347 | 108.5 ± 5.158 | -18.25 ± 6.442 |
| ExN-ChR2 | 38.31 ± 4.086 | 120.9 ± 8.758 | -22.95 ± 8.541 |
| p-value | p= 0.8832 | p= 0.2057 | p= 0.6574 |

Data were presented as mean ± SEM. p > 0.05, unpaired two-tailed Student’s t-test. CTL, n = 20 cells from 9 mice; ExN-ChR2, n = 14 cells from 9 mice.

### Preservation of hippocampal structural connectivity and synapse density

To determine whether the dysregulation of synaptic plasticity and transmission was caused by cellular degeneration, we next assessed changes in the density of pyramidal excitatory neurons within the hippocampus of kindled ExN-ChR2 mice. Unlike other conventional TLE animal models exhibiting widespread necrotic cell death (Lévesque and Avoli, 2013; Mello et al., 1993; Thom, 2009), quantitative histological analysis of the hippocampal CA1 pyramidal layer revealed no significant difference in both the total cell number and overall thickness of the CA1 pyramidal layer (SP) in ExN-ChR2 mice (Fig. 4a-c). These results indicate that behavioral and neurophysiological phenotypes of our optogenetic TLE model do not originate from excitotoxic neurodegeneration.

**Figure 4.**
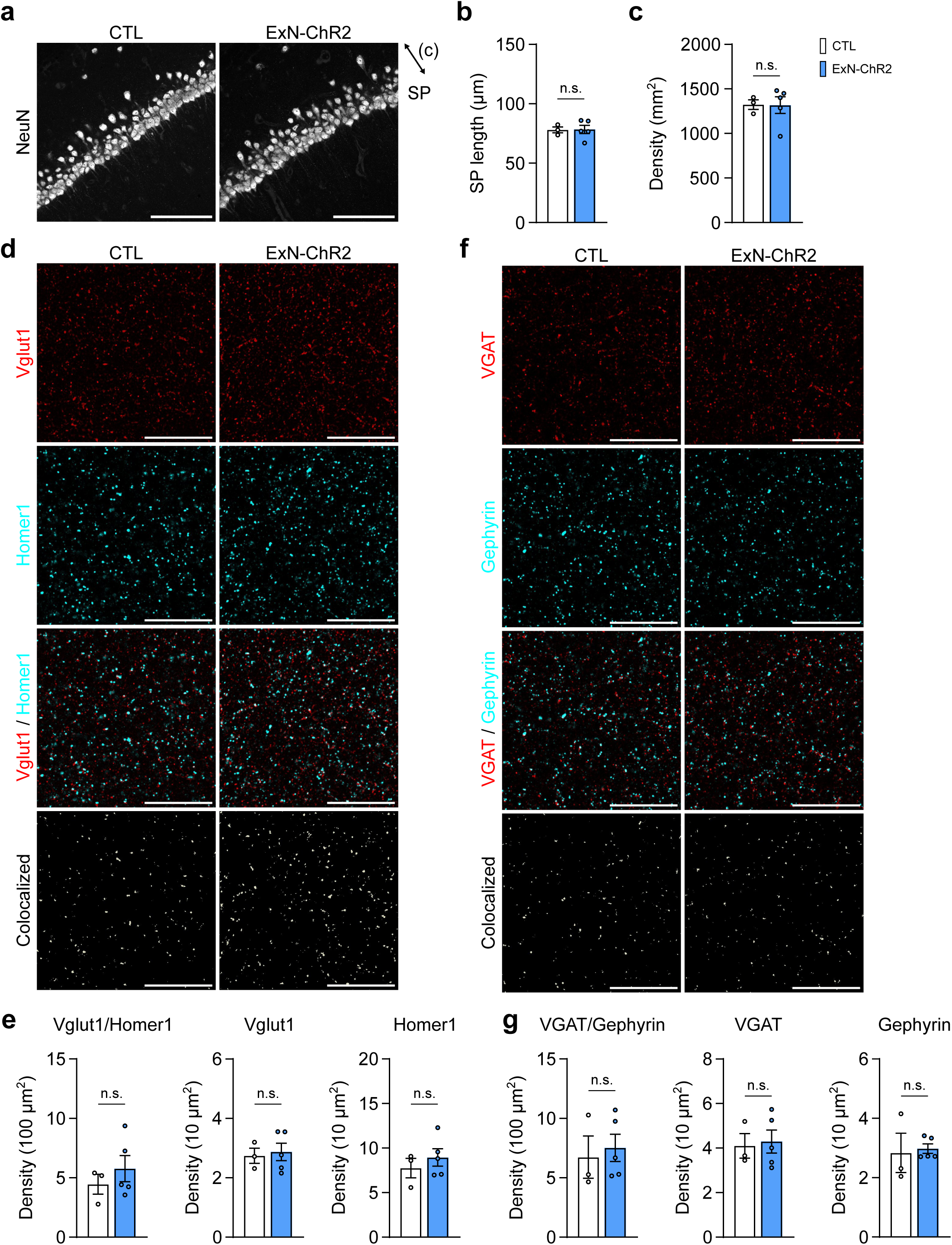
ExN-ChR2 mice show neither neuronal loss nor changes in synaptic puncta density in the CA1 region. a. Representative NeuN immunostaining of the hippocampal CA1 pyramidal layer. The double-headed arrow indicates the thickness of the stratum pyramidale (SP). Scale bar, 100 µm. b. Average SP thickness. P > 0.05. c. Average density of NeuN-positive cells in the pyramidal layer. p > 0.05. d. Representative images of Vglut1 (red), Homer1 (cyan) staining results, the merged image (Vglut1/Homer1), and the extracted colocalized puncta. Scale bars, 50 µm. e. Average density of colocalized Vglut1/Homer1 (per 100 µm²), Vglut1, and Homer1 puncta (per 10 µm²). p > 0.05. f. Representative images of VGAT (red), Gephyrin (cyan), the merged image (VGAT/Gephyrin), and the extracted colocalized puncta. Scale bars, 50 µm. g. Average density of colocalized VGAT/Gephyrin (per 100 µm²), VGAT, and Gephyrin puncta (per 10 µm²). p > 0.05. All data from CTL, n = 3; ExN-ChR2, n = 5 mice. All comparisons were tested using unpaired Welch’s t-test. All data are presented as mean ± SEM.

Alterations in synaptic transmission and plasticity could be evoked by structural remodeling at the synaptic level (Holtmaat and Svoboda, 2009; Matsuzaki et al., 2004, 2001). To test this possibility, we quantified excitatory and inhibitory synapse densities within the hippocampal CA1 stratum radiatum (SR). Synaptic densities were evaluated by analyzing markers including presynaptic vesicular glutamate transporter 1 (Vglut1) and postsynaptic homer scaffold protein 1 (Homer1) for excitatory synapses and vesicular inhibitory amino acid transporter (VGAT) and gephyrin for inhibitory synapses (Fig. 4d-g). Our results showed no significant differences either in excitatory or inhibitory synapse density between groups (Fig. 4d-g). These findings indicate that abnormal synaptic plasticity and transmission in kindled ExN-ChR2 mice are independent of structural synapse loss or remodeling.

### Abnormal astrogliosis in kindled ExN-ChR2 mice

Homeostatic plasticity could be induced by several mechanisms including reactive glia-derived factors such as tumor necrosis factor alpha (TNF-α) (Leonoudakis et al., 2004; Stellwagen and Malenka, 2006). Moreover, increased gliosis is a well-known characteristic of TLE (Vezzani et al., 2022). We thus evaluated morphological changes in astrocytes and microglia upon optogenetic kindling.

In TLE pathogenesis, microglial activation typically predominates during the initial phase, whereas reactive astrocytes drive chronic pathogenic progression (Sano et al., 2021). Consistent with this progressive glial transition, we found robust reactive astrogliosis in the hippocampus of kindled ExN-ChR2 mice, evidenced by upregulated glial fibrillary acidic protein (GFAP) levels and process arborization after chronic kindling of ExN-ChR2 (Fig. 5a-e; Supplementary Fig. 4a-f). However, ionized calcium-binding adapter molecule 1 (Iba1) expression levels and microglial morphological complexity (Sholl analysis) remained intact after the chronic kindling phase (Fig. 5f-k). These results suggest that reactive astrocytes during the progression of TLE contribute to the homeostatic downregulation of excitatory synapses.

**Figure 5.**
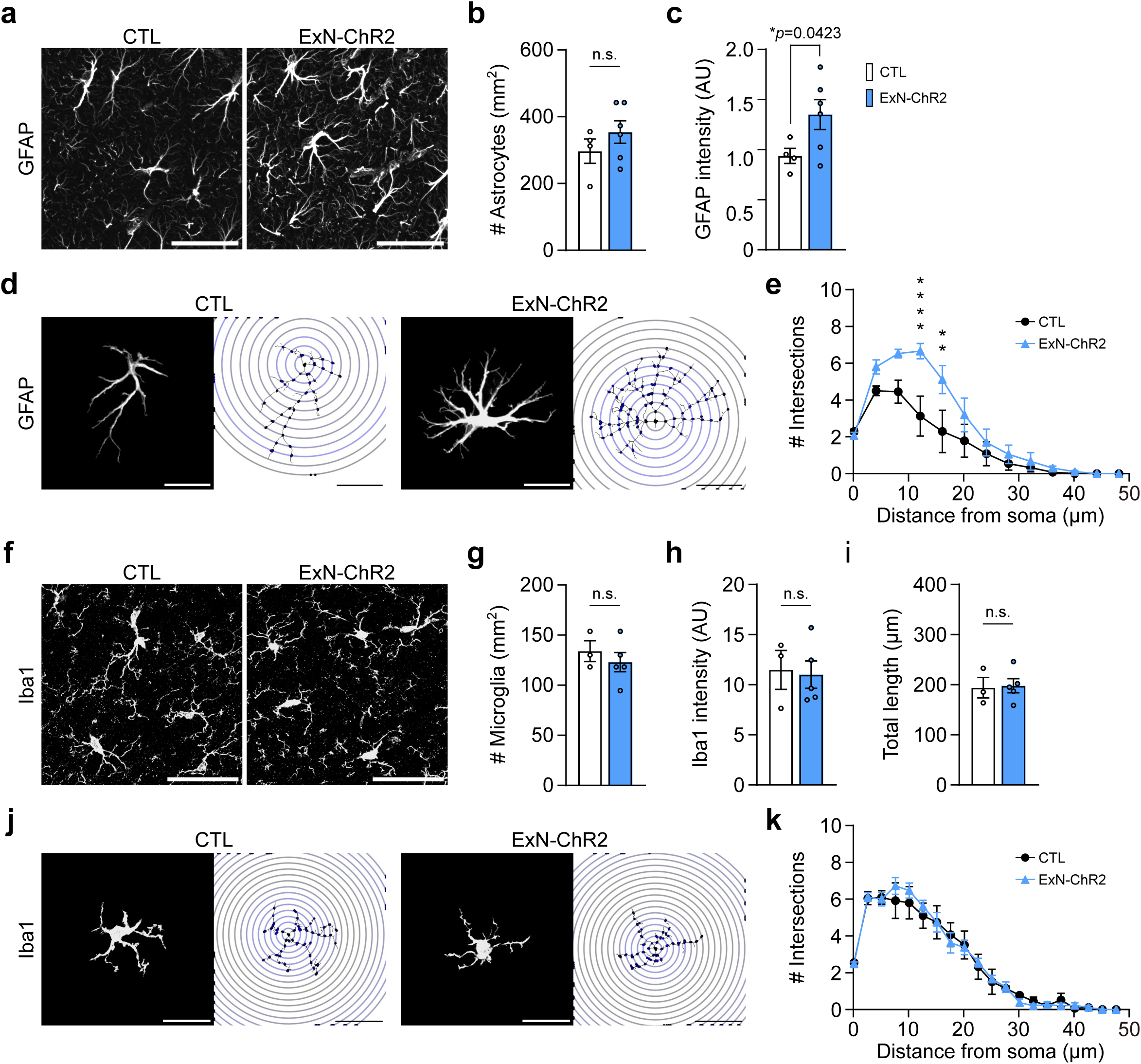
Astrocytic, but not microglial, reactivity in the CA1 region of ExN-ChR2 mice. a. Representative images of GFAP-positive astrocytes in CA1. Scale bar, 50 µm. b. Density of GFAP-positive astrocytes (Number of astrocytes/mm^2^). P > 0.05, unpaired Welch’s t-test. c. Mean GFAP fluorescence intensity. *p = 0.0423, unpaired Welch’s t-test. d. Representative individual GFAP-positive astrocytes (left) and the corresponding digital reconstructions with 4 μm-concentric Sholl rings (right). Scale bars, 20 µm. e. Number of intersections of individual astrocytes. x = 12.15, ****p < 0.0001; x = 16.15, **p = 0.0034, two-way ANOVA with Sidak’s post hoc test. f. Representative images of Iba1-positive microglia in CA1. Scale bars, 50 µm. g. Density of Iba1-positive microglia (Number of microglia/mm^2^). p > 0.05, unpaired Welch’s t-test. h. Mean Iba1 fluorescence intensity. p > 0.05, unpaired Welch’s t-test. i. Total process length of individual microglia. p > 0.05, unpaired Welch’s t-test. j. Representative individual Iba1-positive microglia and the corresponding digital reconstructions with 2.5 μm-concentric Sholl rings (right). Scale bars, 20 µm. k. Number of intersections of individual microglia. p > 0.05, two-way ANOVA. All data are presented as mean ± SEM.

### Upregulation of TMEM230 in the hippocampus of ExN-ChR2 mice

Given that synaptic integrity was structurally maintained, we next investigated molecular candidates that could account for the selective impairment in glutamatergic transmission. We focused on TMEM230, a transmembrane protein involved in intracellular trafficking, and elevated TMEM230 expression is also reported in postmortem human TLE brain (Zhang et al., 2024). Consistent with findings from human study, immunohistochemical analysis revealed a significant increase in TMEM230 immunoreactivity within the hippocampal CA1 region of ExN-ChR2 mice compared to control mice (Fig. 6a, b).

**Figure 6.**
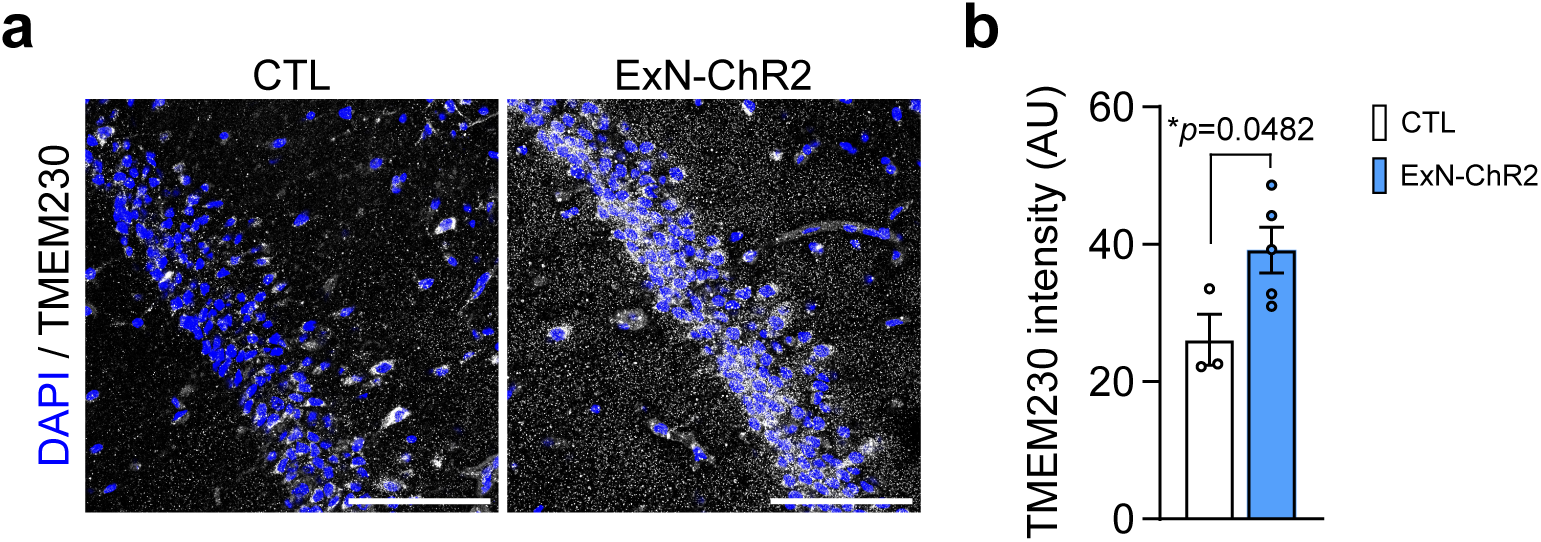
Increased TMEM230 immunoreactivity in the CA1 region of ExN-ChR2 mice. a. Representative images of TMEM230 (white) and DAPI (blue) in the hippocampal CA1 region. Scale bar, 100 µm. b. Mean TMEM230 fluorescence intensity. *p = 0.0482, unpaired Welch’s t-test. CTL, n = 3; ExN-ChR2, n = 5 mice. Data is presented as mean ± SEM.

Given the preferential interaction with presynaptic excitatory machinery, Vglut1, elevated TMEM230 in kindled ExN-ChR2 mice may be selectively involved in homeostatic plasticity of glutamatergic transmission.

## Discussion

In this study, we present an optogenetics-based TLE mouse model. Our optogenetic kindling paradigm could recapitulate the gradually developed TLE features including persistent seizure susceptibility, behavioral comorbidity, and circuit dysfunction. Optogenetics provides an opportunity to model epileptogenesis with cell-type and temporal precision. Repeated excitatory optogenetic stimulation of hippocampal principal neurons can induce progressive, kindling-like seizure development. Crucially, this strategy isolates seizure induction from the confounding neurotoxicity typically observed with chemical convulsants or direct electrical stimulation. Importantly, our lesion/degeneration-free model provides a highly relevant conceptual and translational framework for non-lesional TLE of human patients. In clinical settings, TLE is highly heterogeneous; while a subset of patients presents with classical hippocampal sclerosis marked by severe neurodegeneration, approximately 30 ∼ 40 % of TLE patients exhibit no detectable hippocampal atrophy or neuronal loss on conventional neuroimaging (Vaughan et al., 2016; Wang et al., 2013). Therefore, our model can be useful for studying etiology of TLE patients with non-neurodegenerative phenotypes governed primarily by functional synaptic dysregulation.

Unlike previous optogenetic kindling studies (Tescarollo et al., 2023), the present work offers several novel perspectives in the network and synapse levels during progressive epileptogenesis. Our study provides evidence that chronic focal hyperactivity in hippocampal CA1 excitatory neurons systematically reorganize synaptic efficacy and disrupts E/I balance through homeostatic plasticity (Fig. 3). This result is partially consistent with previous finding that chronic excitation of hippocampal neurons results in homeostatic downregulation of excitatory synapses (Moulin et al., 2019). These homeostatic downregulation of excitatory synapses could be mediated by (1) a postsynaptic α-amino-3-hydroxy-5-methyl-4-isoxazolepropionic acid (AMPA) receptor (AMPAR) internalization or functional desensitization, such as reduced GluA1/GluA2 surface expression (Evers et al., 2010; Goold and Nicoll, 2010) or (2) a selective reduction in basal presynaptic release probability (Pr) within spontaneous vesicle pools, or an increase in functionally silent synapses (e.g., AMPAR-lacking or low-Pr synaptic contacts).

Moreover, additional mechanisms over intrasynaptic mechanisms such as reactive astrocytes and TMEM230 overexpression (Fig. 5 and 6) may contribute to optogenetic kindling-induced homeostatic downregulation. For example, reactive astrocytes can participate in reducing the insertion of AMPA receptors at excitatory synapses by releasing TNF-α (Leonoudakis et al., 2004; Stellwagen and Malenka, 2006) or induce the hyperexcitability by causing the imbalance of ion homeostasis, delayed neurotransmitter clearance, and releasing active gliotransmitters (Álvarez-Ferradas et al., 2015; Ortinski et al., 2010; Shandra et al., 2019; Vezzani et al., 2022). Due to the preferential interaction with excitatory synaptic vesicles, elevated TMEM230 at excitatory synapses may downregulate recycling or docking neurotransmitter-containing vesicles (Zhang et al., 2024). Because such increased astrogliosis and TMEM230 expression are observed from TLE patients (Zhang et al., 2024), our results suggest that homeostatic plasticity-mediated synaptopathy is one of the major drivers of TLE pathology.

Chronic and repetitive hyperactivation by optogenetic kindling is expected to drive circuits into a state of plastic saturation, imposing a ceiling effect that limits further activity-dependent potentiation. Accordingly, we found a severe impairment in CA3-CA1 LTP, a cellular mechanism for associative memory, that is sensitive to homeostatic plasticity mechanisms (Evers et al., 2010; Goold and Nicoll, 2010). This impaired LTP likely contributes to the deficits in hippocampus-dependent cognitive functions (Fig. 2).

In summary, our optogenetic model shows that chronic, cell-type-specific hyperactivation of hippocampal CA1 neurons reproduces the full behavioral and electrophysiological spectrum of TLE without neuronal degeneration. This study provides novel evidence for homeostatic plasticity-induced synaptopathy as a main player in TLE progress.

## Acknowledgements

This work was supported by the KBRI basic research program (26-BR-01-03 and 26-BR-ISD-01) and the National Research Foundation of Korea(NRF) grant funded by the Korean government (MSIT) RS-2023-00265524.

## References

Feigin, V. L. et al. Global, regional, and national burden of epilepsy, 1990-2021: a systematic analysis for the Global Burden of Disease Study 2021. Lancet Public Health 10, e203–e227 (2025).

Téllez-Zenteno, J. F. & Hernández-Ronquillo, L. A Review of the Epidemiology of Temporal Lobe Epilepsy. Epilepsy Res. Treat. 2012, 630853 (2012).

Chen, Z., Brodie, M. J., Liew, D. & Kwan, P. Treatment Outcomes in Patients With Newly Diagnosed Epilepsy Treated With Established and New Antiepileptic Drugs: A 30-Year Longitudinal Cohort Study. JAMA Neurol. 75, 279–286 (2018).

Lévesque, M., Avoli, M. & Bernard, C. Animal models of temporal lobe epilepsy following systemic chemoconvulsant administration. J. Neurosci. Methods 260, 45–52 (2016).

Buckmaster, P. S. & Dudek, F. E. Network Properties of the Dentate Gyrus in Epileptic Rats With Hilar Neuron Loss and Granule Cell Axon Reorganization. J. Neurophysiol. 77, 2685–2696 (1997).

Goldberg, E. M. & Coulter, D. A. Mechanisms of epileptogenesis: a convergence on neural circuit dysfunction. Nat. Rev. Neurosci. 14, 337–349 (2013).

Curia, G., Longo, D., Biagini, G., Jones, R. S. G. & Avoli, M. The pilocarpine model of temporal lobe epilepsy. J. Neurosci. Methods 172, 143–157 (2008).

Wuarin, J.-P. & Dudek, F. E. Electrographic Seizures and New Recurrent Excitatory Circuits in the Dentate Gyrus of Hippocampal Slices from Kainate-Treated Epileptic Rats. The Journal of Neuroscience 16, 4438 (1996).

Mello, L. E. A. M. et al. Circuit Mechanisms of Seizures in the Pilocarpine Model of Chronic Epilepsy: Cell Loss and Mossy Fiber Sprouting. Epilepsia 34, 985–995 (1993).

Lévesque, M. & Avoli, M. The kainic acid model of temporal lobe epilepsy. Neurosci. Biobehav. Rev. 37, 2887–2899 (2013).

Thom, M. Hippocampal sclerosis: Progress since sommer. Brain Pathology vol. 19 565–572 Preprint at 10.1111/j.1750-3639.2008.00201.x (2009).

Vezzani, A. et al. Astrocytes in the initiation and progression of epilepsy. Nat. Rev. Neurol. 18, 707–722 (2022).

Patel, D. C., Tewari, B. P., Chaunsali, L. & Sontheimer, H. Neuron–glia interactions in the pathophysiology of epilepsy. Nat. Rev. Neurosci. 20, 282–297 (2019).

Vaughan, D. N., Rayner, G., Tailby, C. & Jackson, G. D. MRI-negative temporal lobe epilepsy. Neurology 87, 1934–1942 (2016).

Wang, Z. I. et al. The pathology of magnetic-resonance-imaging-negative epilepsy. Modern Pathology 26, 1051–1058 (2013).

Blümcke, I. et al. International consensus classification of hippocampal sclerosis in temporal lobe epilepsy: A Task Force report from the ILAE Commission on Diagnostic Methods. Epilepsia 54, 1315–1329 (2013).

Tescarollo, F. C., Valdivia, D., Chen, S. & Sun, H. Unilateral optogenetic kindling of hippocampus leads to more severe impairments of the inhibitory signaling in the contralateral hippocampus. Front. Mol. Neurosci. Volume 16-2023, (2023).

Buckmaster, P. S., Reyes, B., Kahn, T. & Wyeth, M. Ventral Hippocampal Formation Is the Primary Epileptogenic Zone in a Rat Model of Temporal Lobe Epilepsy. The Journal of Neuroscience 42, 7482 (2022).

Zhang, H. et al. Transmembrane protein modulates seizure in epilepsy: evidence from temporal lobe epilepsy patients and mouse models. Exp. Anim. 73, 162–174 (2024).

Vossler, D. G. et al. Intracranial EEG in Temporal Lobe Epilepsy: Location of Seizure Onset Relates to Degree of Hippocampal Pathology. Epilepsia 45, 497–503 (2004).

Bragin, A., Engel, J. & Staba, R. High-frequency oscillations in epileptic brain. Curr. Opin. Neurol. 23, 151–156 (2010).

Jabran, Y. et al. Temporal and Spatial Dynamics of Different Interictal Epileptic Discharges: A Time-Frequency EEG Approach in Pediatric Focal Refractory Epilepsy. Front. Neurol. Volume 11-2020, (2020).

Fiest, K. M. et al. Depression in epilepsy. Neurology 80, 590–599 (2013).

Keezer, M. R., Sisodiya, S. M. & Sander, J. W. Comorbidities of epilepsy: current concepts and future perspectives. Lancet Neurol. 15, 106–115 (2016).

Hermann, B. P. et al. Cognitive prognosis in chronic temporal lobe epilepsy. Ann. Neurol. 60, 80–87 (2006).

Vinti, V. et al. Temporal Lobe Epilepsy and Psychiatric Comorbidity. Front. Neurol. Volume 12-2021, (2021).

Tao, C. et al. Excitation-inhibition imbalance in medial preoptic area circuits underlies chronic stress-induced depression-like states. Nat. Commun. 15, 8575 (2024).

Li, J. et al. Lack of interferon regulatory factor 3 leads to anxiety/depression-like behaviors through disrupting the balance of neuronal excitation and inhibition in mice. Genes Dis. 10, 1062–1074 (2023).

Matsuzaki, M. et al. Dendritic spine geometry is critical for AMPA receptor expression in hippocampal CA1 pyramidal neurons. Nat. Neurosci. 4, 1086–1092 (2001).

Matsuzaki, M., Honkura, N., Ellis-Davies, G. C. R. & Kasai, H. Structural basis of long-term potentiation in single dendritic spines. Nature 429, 761–766 (2004).

Holtmaat, A. & Svoboda, K. Experience-dependent structural synaptic plasticity in the mammalian brain. Nat. Rev. Neurosci. 10, 647–658 (2009).

Leonoudakis, D., Braithwaite, S. P., Beattie, M. S. & Beattie, E. C. TNFα-induced AMPA-receptor trafficking in CNS neurons; relevance to excitotoxicity? Neuron Glia Biol. 1, 263–273 (2004).

Stellwagen, D. & Malenka, R. C. Synaptic scaling mediated by glial TNF-α. Nature 440, 1054–1059 (2006).

Sano, F., et al. Reactive astrocyte-driven epileptogenesis is induced by microglia initially activated following status epilepticus. JCI Insight 6, (2021).

Moulin, T. C. et al. Chronic in vivo optogenetic stimulation modulates neuronal excitability, spine morphology, and Hebbian plasticity in the mouse hippocampus. Hippocampus 29, 755–761 (2019).

Goold, C. P. & Nicoll, R. A. Single-Cell Optogenetic Excitation Drives Homeostatic Synaptic Depression. Neuron 68, 512–528 (2010).

Evers, D. M. et al. Plk2 attachment to NSF induces homeostatic removal of GluA2 during chronic overexcitation. Nat. Neurosci. 13, 1199–1207 (2010).

Ortinski, P. I. et al. Selective induction of astrocytic gliosis generates deficits in neuronal inhibition. Nat. Neurosci. 13, 584–591 (2010).

Shandra, O. et al. Repetitive Diffuse Mild Traumatic Brain Injury Causes an Atypical Astrocyte Response and Spontaneous Recurrent Seizures. The Journal of Neuroscience 39, 1944 (2019).

Álvarez-Ferradas, C. et al. Enhanced astroglial Ca2+ signaling increases excitatory synaptic strength in the epileptic brain. Glia 63, 1507–1521 (2015).

