## supplementary figure and legend for "Optogenetic Kindling Model Reveals Functional Synaptopathy as a Driver of Temporal Lobe Epilepsy and Associated Comorbidities"

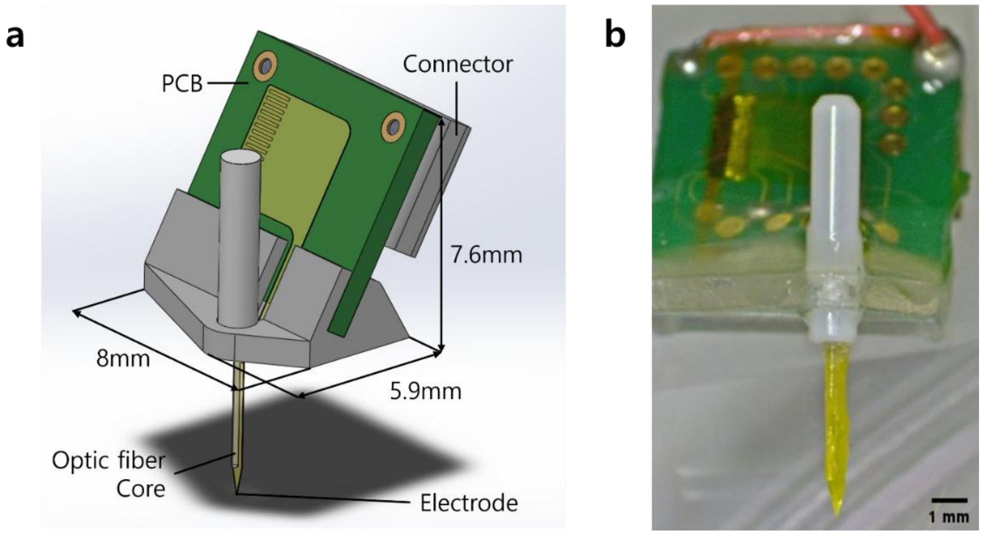


**Supplementary Figure 1. Optrode, optic fiber-neural electrode integrated system.**

a. 3D schematic illustration of the optrode. b. Photograph of the fabricated integrated system. The fabricated electrode array was connected to a printed circuit board (PCB) via anisotropic conductive film (ACF) bonding, and a 16-channel connector (Omnetics) was soldered onto the PCB. The PCB-electrode assembly and an optical fiber were integrated into a 3D-printed housing, where the electrode shank and the optical fiber were bonded together using polyethylene glycol (PEG).

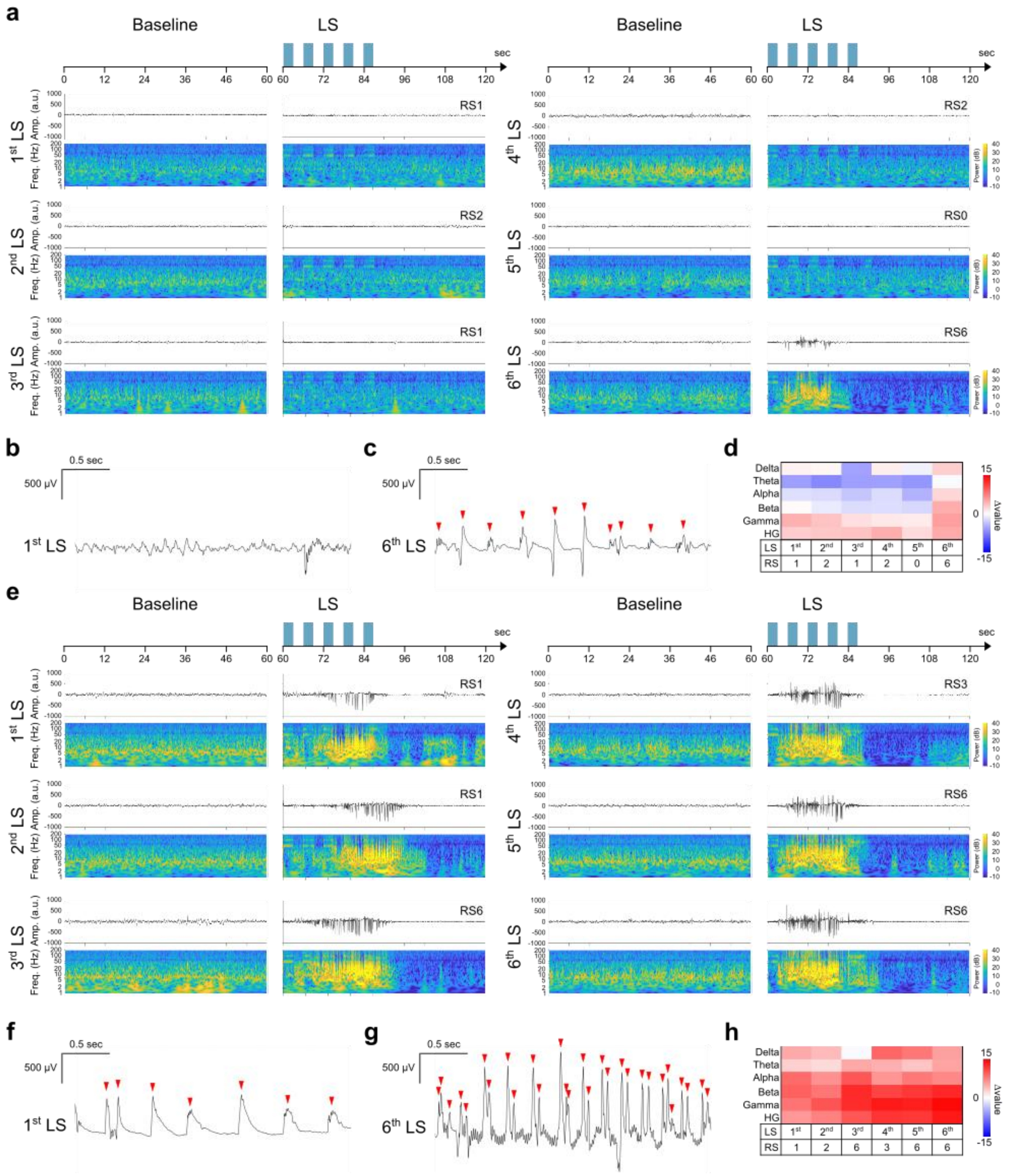


**Supplementary Figure 2. LFP recordings during the first and second re-kindling**

a-d. LFP recordings during the first re-kindling. a. LFP traces analyzed 1 min before and after light stimulation (LS) onset. b-c. Expanded 3-s LFP traces showing individual spikes (red arrows) during 1$^{st}$ (b) and 6$^{th}$ (c) LS session. d. Power spectral density (PSD) of LFP, expressed in dB relative to baseline with Racine score (RS). e-h. LFP recordings during the second re-kindling. e. LFP traces analyzed 1 min before and after LS onset. f-g. Expanded 3-s LFP traces showing individual spikes (red arrows) during 1$^{st}$ (f) and 6$^{th}$ (g) LS session. h. Power spectral density (PSD) of LFP, expressed in dB relative to baseline with RS.

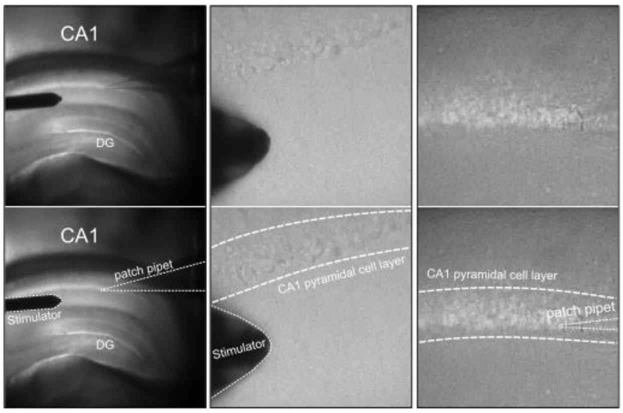


**Supplementary Figure 3. Experimental setup for evoked EPSC and IPSC**

Low-magnification overview (left) showing the CA1 and dentate gyrus (DG) regions, with corresponding high-magnification images focusing on the stimulator placement (middle) and the patch pipet targeted at the CA1 pyramidal cell layer (right). Dashed lines delineate the contours of the stimulator, the boundaries of the CA1 cell layer, and the trajectory of the patch pipet.

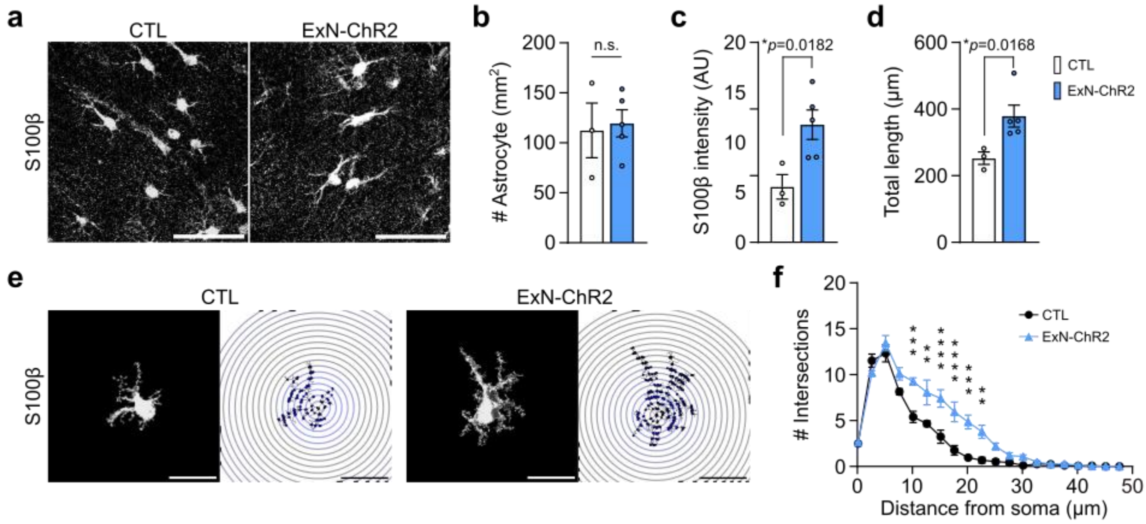


**Supplementary Figure 4. Astrocytic reactivity in the CA1 region of ExN-ChR2 mice.**

a. Representative images of S100β-positive astrocyte in CA1. Scale bars, 50 µm. g. Density of S100β-positive astrocyte (Number of astrocyte/mm$^2$). p> 0.05, unpaired Welch's t-test. h. Mean S100β fluorescence intensity. *p= 0.0182, unpaired Welch's t-test. i. Total process length of individual astrocyte. *p= 0.0168, unpaired Welch's t-test. j. Representative individual S100β-positive astrocyte and the corresponding digital reconstructions with 2.5 µm-concentric Sholl rings (right). Scale bars, 20 µm. k. Number of intersections of individual astrocyte. x= 10.14, ***p= 0.0002; x= 12.64, **p= 0.0015; x= 15.14, ****p= 0.000033; x= 17.64, ****p= 0.000031; x= 20.14, ***p= 0.0001; x= 22.64, **p= 0.0051, two-way ANOVA with Sidak's post hoc test. All data are presented as mean ± SEM.
